# Geometric morphometrics of nested symmetries: Hierarchical INTER- AND INTRA-INDIVIDUAL VARIATION IN BIOLOGICAL SHAPES

**DOI:** 10.1101/306712

**Authors:** Yoland Savriama, Sylvain Gerber

**Affiliations:** Institute of Biotechnology, PO Box 56 (Viikinkaari 5), FIN-00014 University of Helsinki, Helsinki, Finland; Institut de Systématique, Évolution, Biodiversité ISYEB - UMR 7205 - MNHN CNRS UPMC EPHE, Muséum national d’Histoire naturelle, Sorbonne Universités, 57 rue Cuvier, CP 50, 75005 Paris, France

**Keywords:** complex symmetry, developmental instability, fluctuating asymmetry, nested design, Procrustes ANOVA, shape analysis

## Abstract

Symmetry is a pervasive feature of organismal shape and the focus of a large body of research in Biology. Here, we consider complex patterns of symmetry where a phenotype exhibits a hierarchically structured combination of symmetries. We extend the Procrustes ANOVA for the analysis of nested symmetries and the decomposition of the overall morphological variation into components of symmetry (among-individual variation) and asymmetry (directional and fluctuating asymmetry). We illustrate its use with the Aristotle’s lantern, the masticatory apparatus of ‘regular’ sea urchins, a complex organ displaying bilateral symmetry nested within five-fold rotational symmetry. Our results highlight the importance of characterising the full symmetry of a structure with nested symmetries. Higher order rotational symmetry appears strongly constrained and developmentally stable compared to lower level bilateral symmetry. This contrast between higher and lower levels of asymmetry is discussed in relation to the spatial pattern of the lantern morphogenesis. This extended framework is applicable to any biological object exhibiting nested symmetries, regardless of their type (e.g., bilateral, rotational, translational). Such cases are extremely widespread in animals and plants, from arthropod segmentation to angiosperm inflorescence and corolla shape. The method therefore widens the research scope on developmental instability, canalization, developmental modularity and morphological integration.

## Introduction

The anatomical organisation of almost all organisms appear to follow precise patterns of symmetry in which body parts are repeated and geometrically arranged according to specific positions and orientations. As an important feature of morphological phenotypes, symmetry is at the core of numerous research programs in evolutionary and developmental biology, such as the origin of symmetry in the corolla of flowers (e.g. Citerne et al. 2010), the mechanisms of phyllotaxis and branching in plants (e.g. Reinhardt et al. 2003), and the patterns of segmentation and serial homology (e.g. Fusco and Minelli 2013). Deviation from symmetry, or asymmetry, is also critical to our understanding of the origin of phenotypic variation. The measurement of directional asymmetry, the population average systematic deviation from perfect symmetry, and of fluctuating asymmetry, the small random deviations from perfect symmetry or around the average directional asymmetry, underlies research on developmental instability, canalization, morphological integration, and developmental modularity (Klingenberg 2003; Møller and Swaddle 1997; Palmer and Strobeck 1986; Polak 2003).

The study of biological forms has greatly benefited from the advent of geometric morphometric methods (GMMs), a powerful array of mathematical and statistical tools for quantifying and analysing shape and shape variation. Geometric morphometric frameworks have also been devised for the study of symmetry and asymmetry in bilaterally symmetric structures, and these frameworks have been recently generalized to all types of symmetry (Klingenberg et al. 2002; Mardia et al. 2000; Savriama and Klingenberg 2011). Morphometric analyses of symmetric structures allow the partitioning of the total variation into components of symmetric and asymmetric variation. A two-way mixed model Procrustes ANOVA further decomposes this total variation into biologically meaningful components. Its design includes the main effect ‘individual’ describing variation among individuals, the main effect ‘side’ standing for directional asymmetry (DA), the ‘individual × side’ interaction standing for fluctuating asymmetry (FA), and measurement error if two or more replicates have been taken (Klingenberg and McIntyre 1998).

More complex, composite types of symmetry can be found in nature. They correspond to cases where a phenotype displays a hierarchically structured combination of symmetries. The vertebral column is for instance composed of bilaterally symmetric vertebrae that are themselves arranged according to translational symmetry along the body axis. This same type of nested symmetries is exhibited in arthropod segmentation (e.g. Fusco and Minelli 2013). Plants also show nested symmetries. In angiosperms, the tight arrangement of flowers forming a capitulum is such that it generates spiral symmetry, while the flowers themselves have symmetries ranging from bilateral symmetry (zygomorphy), left-right asymmetry, to rotational symmetry as their distance from the centre of the capitulum increases (e.g. Berger et al. 2016; Carlson et al. 2011).

How should the total variation be decomposed in a sample of phenotypes with nested symmetries? Central to the study of symmetric phenotypes is the identification of the repeated unit underlying the symmetry of the phenotype considered. In the case of ambiguous symmetry where distinct symmetry groups can account for the symmetric pattern observed, developmental hypotheses can help identify the unit and the symmetry group involved, and reject competing alternatives suggested by contingent by-products of its symmetry transformations (e.g. corals, Savriama and Klingenberg 2011). For nested symmetries, the key is to consider the repeated unit at the higher level of symmetric organisation as being itself a product of symmetry transformations applied to a lower level unit. Theoretically, each of these levels can be expected to display among-individual variation in symmetric shape and within-individual deviations from perfect symmetry, but little is known about their possible patterns. Are these nested levels of symmetry equally sensitive to developmental perturbations? If not, how are patterns of developmental stability structured across levels and how well do they align with patterns of individual variation?

Here, we extend the statistical design of Procrustes ANOVA to analyse symmetric and asymmetric variations in organisms exhibiting hierarchically organised symmetries. We illustrate this approach with a small dataset of Aristotle’s lanterns, the masticatory apparatus of ‘regular’ sea urchins, which is considered as a key character underlying the adaptive radiation and evolutionary success of sea-urchins (David et al. 2009; de Ridder and Lawrence 1982; Mooi 1990). The lantern is a complex ensemble of calcitic skeletal pieces that all show bilateral symmetry and are themselves nested within the more complex five-fold rotational symmetry typifying the echinoderm body plan.

We consider three possible analyses of symmetry for this ambiguous case study based on three hypotheses regarding which fundamental repeated unit generates the type of symmetry of biological interest: the rotational symmetry of the lantern, the bilateral symmetry of the pyramids, and the bilateral symmetry nested within the rotational symmetry. For each case, we carry out Procrustes analysis and statistical decomposition of the symmetric and asymmetric components of variation in order to unravel and display patterns of interindividual variation, directional asymmetry, and fluctuating asymmetry. We show that the full model of nested symmetries outperforms the alternative models and allows testing the significance of rotational and bilateral asymmetries considered jointly and according to their hierarchical layout. The results indicate contrasted degrees of rotational and bilateral asymmetries, which we discuss in the context of differential developmental stability and functional constraints acting at these two levels.

## Geometric morphometrics of symmetric structures

Bilateral symmetry is the simplest type of symmetry and is relatively widespread in biological systems. Other common but more complex types of symmetry include rotational symmetry (e.g. the corona of ‘regular’ sea urchins), translational symmetry (e.g. the segments of arthropods) or combinations of them (e.g. the spiral symmetry in Molluscan shells). Following Mardia (2000), all these biological instances of symmetry can be classified into two categories of symmetry depending on the features of the biological structure considered: matching symmetry and object symmetry. We now briefly review their general treatment in geometric morphometrics before elaborating the case of nested symmetries.

### Geometric morphometrics of matching symmetry

In the most general case, matching symmetry refers to the case where a structure is composed of a series of *k* repeated units that are physically disconnected and geometrically organised with respect to the symmetry transformation characterising the structure (e.g., reflectional symmetry of the butterfly wings, rotational symmetry of petals in some flowers). For its analysis, a common configuration of homologous landmarks is defined for the repeated unit and is recorded for each of the repeats. If necessary, the configurations are adjusted to place them in a comparable orientation. For instance, in the case of bilateral symmetry, one of the *x*, *y* or *z* coordinates of one of the two sides is multiplied by −1 so that the left and right sides can be meaningfully superimposed. All the configurations are then superimposed by a single Generalized Procrustes Analysis (GPA) which extracts shape variation by filtering out the effects of scale, orientation and translation (e.g. Dryden and Mardia 1998). A resulting mean shape (consensus) is simultaneously estimated and the shape variation around it can be decomposed into components of symmetric and asymmetric variation. Symmetric variation (i.e. variation among individuals) is calculated as the differences among the averages of all parts, and asymmetry (i.e. variation within individuals) is measured as the deviations from the respective individual average parts. Since a separate landmark configuration is recorded for each of the *k* units, components of symmetric and asymmetric variation can also be calculated for size (centroid size).

A two-way mixed model ANOVA with individuals and parts as the two factors is used to partition the total variation into components of interest (Klingenberg and McIntyre 1998; Leamy 1984; Palmer and Strobeck 1986). The main effect of ‘individual’ reflects the variation among individuals (symmetric component), the main effect ‘part’ represents directional asymmetry (the average deviation among parts), and the ‘individual × part’ interaction represents fluctuating asymmetry (small random deviation from the average asymmetry among parts). Measurement error due to imaging and digitizing can be assessed if replicate measurements have been taken.

### Geometric morphometries of object symmetry

Object symmetry refers to symmetric structures for which the symmetry operators (centre, axis or plane) belong to the structure, which is therefore symmetric as a whole (e.g., reflectional symmetry of the human skull, rotational symmetry of a coral polyp). The structure is separable into *k* connected parts. The analysis of structures with object symmetry considers the variation among parts as in matching symmetry, but with additional information about the way the *k* parts are physically connected to each other. Methodologically, a unique configuration of landmarks is considered for the entire structure. For each individual configuration of landmarks, *n* copies are generated, where *n* is the number of transformations that characterises the symmetry group of the structure (Klingenberg et al. 2002; Mardia et al. 2000; Savriama and Klingenberg 2011). For instance, the symmetry group of bilateral symmetry includes two such transformations: identity and reflection with respect to a symmetry plane. These *n* copies are then transformed according to their respective transformation and their landmarks appropriately relabelled. The relabelling procedure simply consists in mutually swapping the labels of the landmarks that are images of each other with respect to the symmetry transformation considered (note that shared landmarks - i.e., located on the symmetry operators - map onto themselves). Thereafter, a GPA is performed on the full dataset which contains all original configurations and their respective transformed and relabelled copies. The resulting consensus is perfectly symmetric with respect to the symmetry group. As for matching symmetry, variation around this average shape can be decomposed into a component of symmetric shape variation (differences among the averages of the original and appropriately transformed and relabelled copies) and a component of asymmetry (within-individual variation, differences between the original and transformed relabelled copies). However, unlike the case of matching symmetry, these components occupy separate subspaces of the shape tangent space that are orthogonal and complementary to each other (Kent and Mardia 2001; Kolamunnage and Kent 2003; Kolamunnage and Kent 2005; Mardia et al. 2000). This particular feature of the shape analysis of object symmetry has specific implications for the statistical testing of ANOVA effects. The same two-way mixed model Procrustes ANOVA still applies, but the main effect of ‘part’ is replaced by the main effect of ‘transformation’ and the calculation of *F*-ratios needs considering the mean squares of the appropriate tangent subspace (Klingenberg et al. 2002; Savriama and Klingenberg 2011). Since there is a unique configuration for the whole structure, there is no asymmetry in size to consider.

## The case of combined symmetries: the Aristotle’s lantern as an example

The Aristotle’s lantern is the masticatory apparatus of ‘regular’ sea urchins. It consists of a set of 40 calcitic skeletal elements activated by a series of complex muscles (Candia Carnevali et al. 1993; Cavey and Märkel 1994; Hyman 1955). Each of these skeletal elements shows bilateral object symmetry and all of them are arranged according to a rotational matching symmetry of order five (Figure 1a). The largest skeletal pieces are the pyramids that draw their names from their triangular shape. Each pyramid is made of two halves or hemipyramids that are mirror images of each other and physically connected by a strong interradial suture. The interior parts of the hemipyramids have elongated skeletal processes operating as scabbards allowing the teeth to slide through them during growth and feeding. The teeth are reinforced by a high content in magnesium that renders them durable as they scrape, cut, chew food and dig holes into hard rocky substrates (de Ridder and Lawrence 1982). The epiphyses are directly connected at the base of these pyramids and act as their further extensions. They can be fused together along the axis of bilateral symmetry or not depending on the species. Each aboral joint of the pyramids are specifically shaped to allow the fitting of a rectangular element called the rotula, effectively connecting the pyramids together. On top of the rotulae and directly sitting on them are narrow and thin elements named the compasses that trigger the tip of the lantern to move in or out of the corona during feeding (Andrietti et al. 1990). Protractor and retractor muscles connect the lantern to the corona via the perignathic girdle which provides anchors called the apophyses that are themselves symmetric and surround the oral opening (peristome) (Ziegler et al. 2012). Other ligaments such as compass depressors and peristomial membrane also exist and are combined with strong muscles between the hemipyramids to exert strong biting and rasping action. These intricate tissue connections between the perignathic girdle and the lantern confers the latter a wide range of motion (Candia Carnevali et al. 1993).

**Figure 1.**
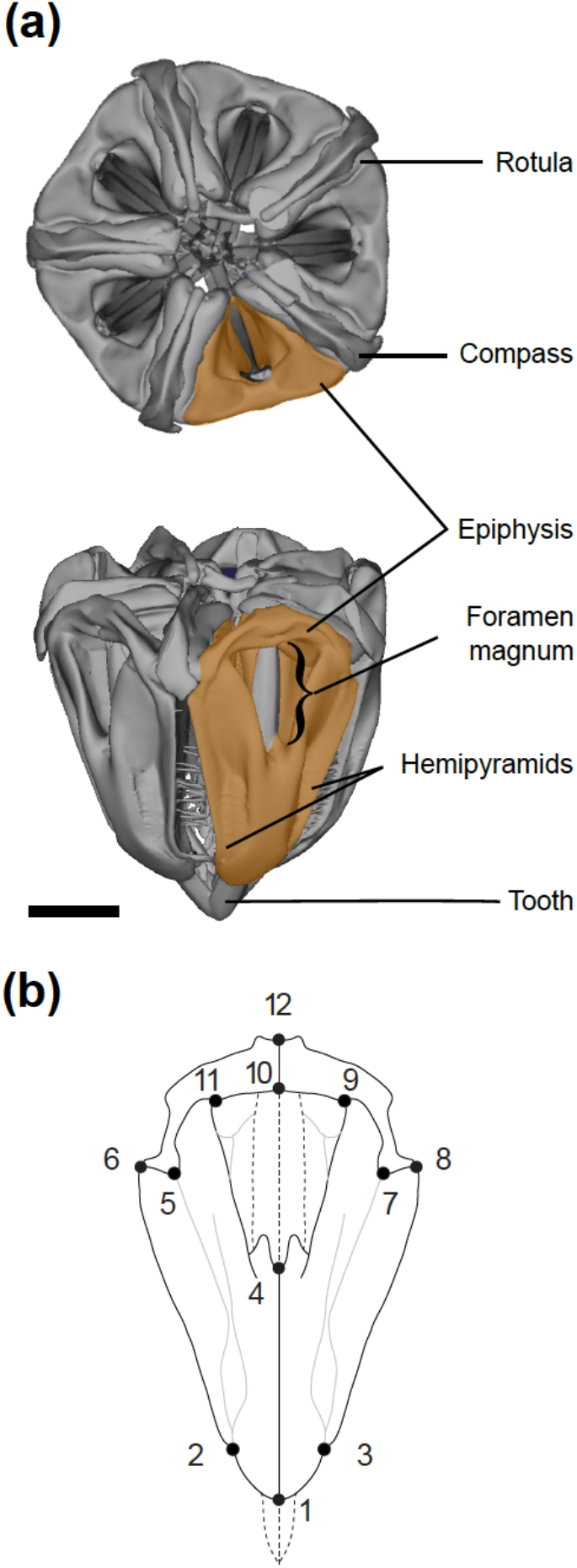
**(a).** CT-Scan of the Aristotle’s lantern of the ‘regular’ sea urchin *P. lividus* showing its nested symmetries. The lower level bilateral object symmetry of the pyramids is nested within the higher level rotational matching symmetry of order five characteristics of echinoderms (see text for details). This hierarchical architecture rests on symmetry transformations applied to sets of hemipyramids and epiphyses (highlighted). Scale bar 5 mm. **(b).** The configuration of homologous anatomical landmarks used to capture the geometry of the pyramids and epiphyses.

## Materials and methods

Here, we analysed a small dataset of pyramids and epiphyses collected from 10 lanterns (50 sets of pyramids plus epiphyses) randomly selected from a larger dataset collected from a population of *Paracentrotus lividus* (Echinodermata, Parechinidae) located in Niolon (Le Rove, France) and that will be fully analysed elsewhere. Prior to any measurement, we recorded the pyramid aligned with the madreporite (a modified genital plate from the corona) in order to effectively orientate the lantern across specimens. This is a critical step to estimate DA in subsequent analyses. Thereafter, all soft tissues were removed by dissolution in hydrochloric acid baths to reveal the skeletal structures of the lantern. A set of 12 landmarks in two dimensions was used to capture the shape of the planar external surface of the pyramids (Figure 1b). The pyramids were photographed in standardized orientation using a Lumenara camera (Model # Infinity2-1C-ACS) mounted to a Zeiss Stemi-2000-C stereomicroscope. Landmarks (Adams and Otárola-Castillo 2013 12) are unpaired landmarks located on the interradial suture between the left and right hemipyramids and epiphyses, i.e., on the plane of bilateral symmetry of the pyramid. Paired landmarks {5, 6, 7, 8} are located at the junction of the hemipyramids and epiphyses. These landmarks also mark the insertion of muscles and other connective soft tissues: Landmark 1 is placed at the bottom-most part of the hemipyramids and connected to the peristomial membrane; landmarks {2, 3} describe the tips of the retractor muscles; landmarks {5, 6, 7, 8, 9, 10, 11, 12} locate the insertion of the protractor muscles, and landmark 4 describes the bottom part of the foramen magnum. To assess measurement error, all measurements were taken twice by the same operator (YS) during separate sessions using tpsDig2 (ver. 2.17, Rohlf 2015). For convenience, we hereafter simply use the term ‘pyramid’ to refer to the combination of the two hemipyramids (true pyramid) with the epiphyses.

We analysed patterns of symmetry and asymmetry at the different levels of symmetric organisation of the Aristotle’s lantern: (1) rotational matching symmetry of order five only, (2) bilateral object symmetry, and (3) bilateral object symmetry nested within rotational matching symmetry of order five (Figure 2). In each case, all transformed and appropriately relabelled copies of the landmark configurations were aligned by partial GPA and orthogonally projected onto the tangent space. All subsequent analyses were performed using the tangent space coordinates. We conducted Procrustes ANOVAs with the statistical design appropriate for each case (see below) and tested for the significance of effects using Goodall’s *F*-test (Goodall 1991). In order to relax the assumption of isotropic shape variation, we also performed MANOVAs with Pillai’s trace as the test statistics (with the use of the generalized inverse for the total sum of squares and cross-products matrix involved in the calculation of Pillai’s trace) (Klingenberg et al. 2002). For both tests, we use parametric and resampling approaches. The degrees of freedom (d.f.) for the parametric tests accounted for the shape dimension of the shape (sub)space considered (conventional d.f. multiplied by the shape dimension). The resampling procedures used 10,000 permutations, with restricted permutations for the main factors, permutations across main factors for the test of interaction after subtraction of their effects, and restricted permutations for the test of nested effects (Anderson and Braak 2003; Edgington and Onghena 2007; Good 2013). Finally, we used Principal Component Analysis (PCA) of the covariance matrices associated with the main effects, interaction terms, and measurement error, to visualize the shape changes associated with each of them. All morphometric and statistical analyses were programmed and carried out in R (Team 2017).

**Figure 2.**
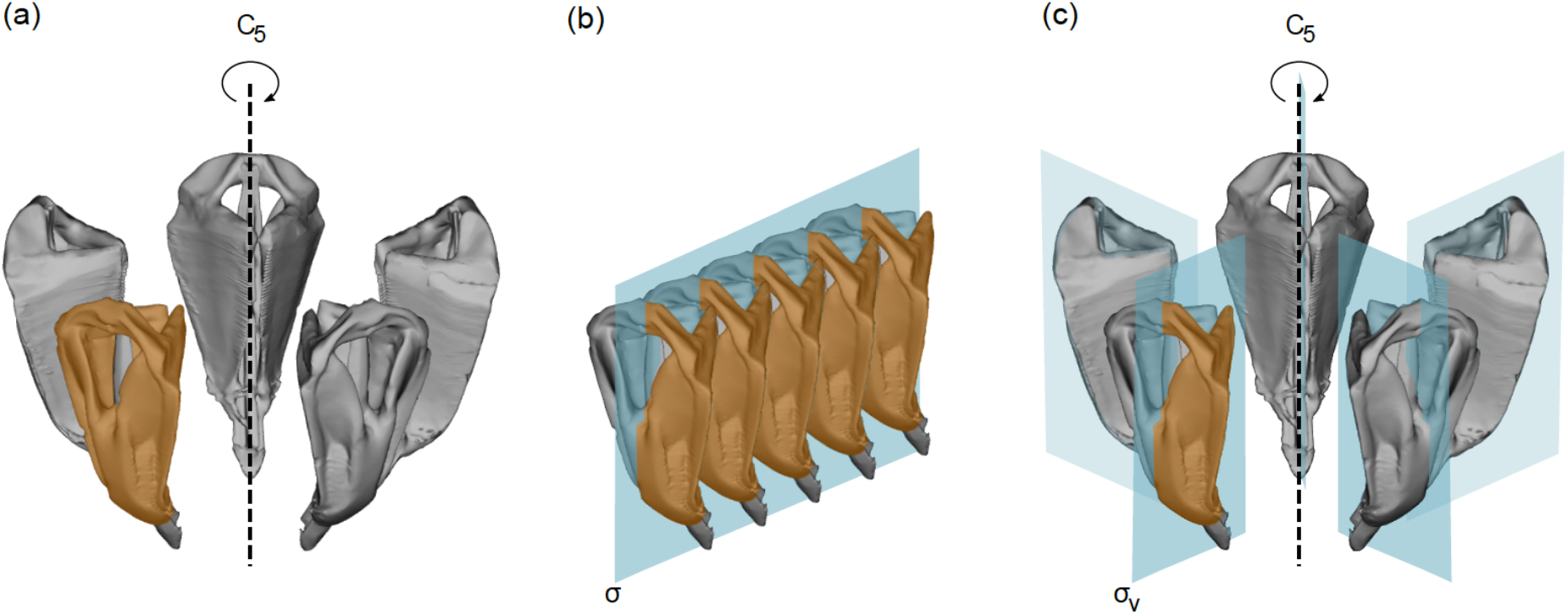
Alternative options for the analysis of the symmetric architecture of the Aristotle’s lantern. (a) rotational matching symmetry of order five only (symmetry group C_5_), (b) bilateral object symmetry only (symmetry group C_1*v*_), and (c) bilateral object symmetry nested within rotational matching symmetry of order five (symmetry group C_5*v*_).

### Analysis 1: Rotational matching symmetry of order five (symmetry group C_5_)

The functional role of the lantern as a masticatory apparatus is ensured by the set of five radially arranged teeth. Their relative positioning, shape, and size are constrained by their firm collagenous attachment to the pyramid allowing them to withstand the mechanical stresses associated with feeding activities (e.g. Birenheide et al. 1996; de Ridder and Lawrence 1982). The adequate closing of the protruding teeth, and thus the relative degree of rotational (a)symmetry among pyramids, is critical for the feeding efficiency (for both soft substrate ingestion and crushing power). In this context, we focus on the five-fold rotational matching shape symmetry of the lantern (symmetry group C_5_) while considering the pyramids as perfectly bilaterally symmetric structures (Figure 2a). For this purpose and prior to the analysis of rotational matching symmetry, each pyramid was treated with the method of object symmetry to extract its symmetric average (the consensus of each pyramid and its reflected copy). The resulting symmetric pyramids are considered as the fundamental units being repeated to generate the rotational symmetry of the lantern. The framework to analyse the rotational symmetry of the lantern is then a natural extension of the approach for studying matching symmetry in bilaterally symmetric structures. The Procrustes ANOVA design is the traditional two-way mixed model with the exception of the ‘side’ effect being replaced by a ‘pyramid’ effect that includes five levels instead of two. It reads as follows (Greek symbols indicate fixed effects and Latin letters stand for random effects):

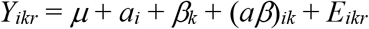

Where *Y_ikr_* is the measurement for lantern *i*, pyramid *k*, replicate *r* (with *i* = 1, 2,…, 10; *k* = 1, 2,…, 5; *r* =1, 2). The parameter *μ* is the grand mean. The main effect *a_i_* is the ‘lantern’ random effect ~ *N (0, σ1)*. The main effect *β_k_* is the ‘pyramid’ effect. It is a fixed effect, since we made a systematic distinction among pyramids by orientating them, representing rotational directional asymmetry (DA). The interaction term (*aβ*)_*ik*_ is a measure of the lantern rotational fluctuating asymmetry (FA) ~ *N (0, σ2). E_ikr_* is measurement error (ME) ~ *N (0, σ3)*. This model extended to *k* repeated units (with *k*>2) was recently used in a study of translational FA in centipedes (Savriama et al. 2016). Size asymmetry can also be analysed since measures of centroid size are available for each pyramid. The same ANOVA design is used for the analysis of size asymmetry but with the conventional degrees of freedom for the calculations of mean squares.

For both size and shape asymmetry analyses, the ANOVAs indicate significant individual variation, taking up most of the overall variation, and significant rotational FA, but non-significant rotational DA (Table 1). The MANOVA, which circumvents the assumption of isotropic variation, gives similar results, and both parametric and non-parametric (permutation) tests provide similar P-values. The absence of rotational DA might appear as a surprise since one could have expected the asymmetric positioning of the oesophagus and digestive tube around the lantern to induce a systematic deviation among the pyramids.

**Table 1.** Procrustes ANOVA, MANOVA, and centroid size ANOVA of the lantern’s fivefold rotational matching symmetry only (analysis 1). Conv. df, conventional degrees of freedom; Shape df, shape degrees of freedom; SS, sum of squares; MS, mean square; F, F-value; P(param), parametric P-value; P(permut), P-value obtained by permutation.

Procrustes ANOVA
| Effect | Conv. df | Shape df | SS | MS | F | P (param) | P (permut) | Pillai's trace | P (param) | P (permut) |
| --- | --- | --- | --- | --- | --- | --- | --- | --- | --- | --- |
| Lantern | 9 | 90 | 0.111301 | 0.001237 | 125.639 | <0.0001 | <0.0001 | 6.8479 | <0.0001 | <0.0001 |
| Pyramid | 4 | 40 | 0.000429 | 0.000011 | 1.090 | 0.33348 | 0.3783 | 1.0868 | 0.3149 | 0.3097 |
| Lantern*Pyramid | 36 | 360 | 0.003544 | 0.000010 | 3.377 | <0.0001 | <0.0001 | 6.8693 | <0.0001 | <0.0001 |
| Measurement | 50 | 500 | 0.001457 | 0.000003 |  |  |  |  |  |  |

Centroid size ANOVA
| Effect | SS | MS | df | F | P (param) | P (permut) |
| --- | --- | --- | --- | --- | --- | --- |
| Lantern | 141.697309 | 15.744145 | 9 | 1353.196 | <0.0001 | <0.0001 |
| Pyramid | 0.037473 | 0.009368 | 4 | 0.805 | 0.53 | 0.3856 |
| Lantern*Pyramid | 0.418852 | 0.011635 | 36 | 7.807 | <0.0001 | <0.0001 |
| Measurement | 0.074512 | 0.001490 | 50 |  |  |  |

The PCA of the covariance matrix for the individual effect reveals that lanterns mostly differ by the relative proportion of their basal part (captured by landmarks 5 to 12), with nearly vertical shifts at the base of the hemipyramids and opposite shifts at the junction between epiphyses (Figure 3). These vertical displacements most likely reflect among-individual variation in the strength of the antagonistic action of the protractor and retractor muscles which have vertical ranges of action. For rotational FA, variation mostly corresponds to the opening of the foramen magnum (PC1), possibly reflecting developmental imprecision during the formation of the interradial suture and interactions with teeth sliding along small bilaterally symmetric hooks within pyramids. PC2 indicates variation in the height-width ratio of the pyramid maybe due to mechanical constraints exerted by the rotulae and the strong interpyramidal muscles located on the external edges of hemipyramids.

**Figure 3.**
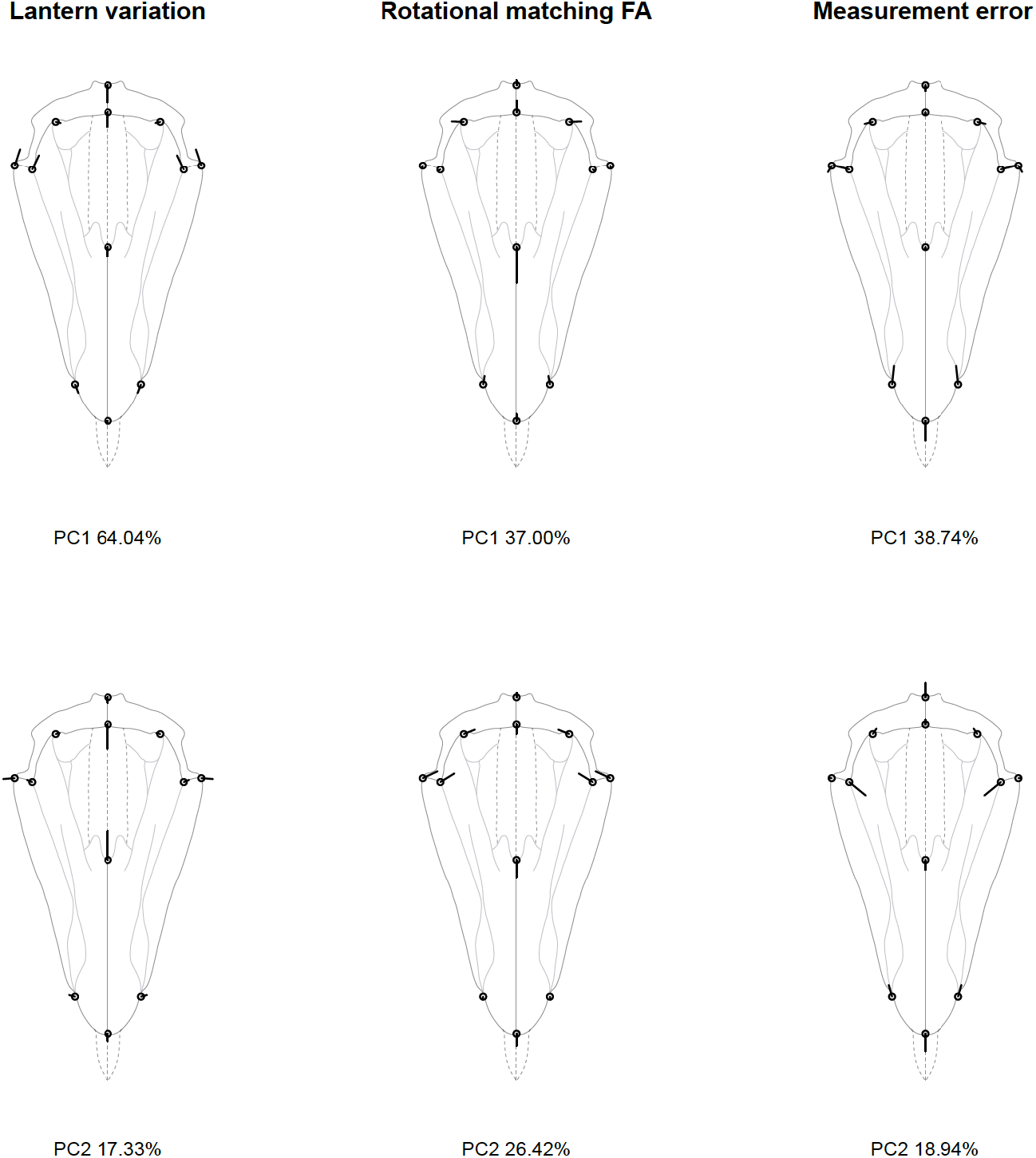
Analysis 1: rotational matching symmetry of order five (C_5_). Principal components describing the patterns of the lantern’s rotational symmetric shape variation, rotational FA, and measurement error shown as lollipop graphs. Open circles represent the consensus configuration and the thick black lines illustrate the magnitude and direction of the vectors of shape change over the first two principal components (PC).

### Analysis 2: Bilateral object symmetry (symmetry group C_1*v*_)

Alongside their diversified habitats and ecological lifestyles, regular sea urchins have a wide range of food types ranging from shelled mollusks to soft substrates (de Ridder and Lawrence 1982). This has led to diverse types of lantern architectures (cidaroid, camarodont, aulodont, strirodont) primarily distinguished by the connectivity of the epiphyses and the depth of the foramen magnum. Despite this substantial diversity of architectures, the bilateral symmetry of the pyramids and epiphyses appears as a pervasive feature of their shapes, suggesting strong functional constraints for the maintenance of symmetry and justifying a particular focus on this level of symmetric organisation. We therefore now consider the bilateral object symmetry of the pyramid, i.e., the differences between the left and right hemipyramids (symmetry group C_1*v*_, Figure 2b). Prior to their analysis we corrected for the individual effect and for individual deviation from rotational symmetry (correction for differences in means), in order to adjust the measurement errors artificially inflated by pooling labelled pyramids from distinct individuals. The total source of variation is decomposed around the mean shape according to the following design (note that the Procrustes ANOVA becomes a two-way model with only fixed main effects):

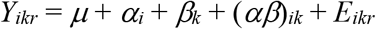

Where *Y_ikr_* is the measurement for pyramid *i*, hemipyramid (or reflection) *k* and replicate *r* (with *i* = 1, 2,…, 5; *k* = 1, 2; *r* =1, 2). The parameter *μ* is the grand mean. The main effect *α_i_* is the ‘pyramid’ fixed effect. The main effect *β_k_* is the ‘hemipyramid’ (or ‘reflection’) effect and measures bilateral DA. The interaction (*αβ*)_*ik*_ is a measure of the pyramid bilateral FA. *E_ikr_* is measurement error (ME) ~ *N(0, σ1)*. Since we are dealing with object symmetry, the tests for the main effects and interaction must consider the error effects from the appropriate symmetric or asymmetric shape subspaces: the symmetric component of measurement error for the ‘pyramid’ effect (whose significance in the MANOVA can be assessed within this fixed model), and the asymmetric component for the ‘hemipyramid’ effect and the interaction term.

The Procrustes ANOVA and the MANOVA both indicate statistically significant ‘pyramid’ effect, ‘hemipyramid’ effect, and ‘pyramid × hemipyramid’ interaction (Table 2). Bilateral object DA is the largest of all effects and expresses systematic differences between left and right hemipyramids. This asymmetry of pyramids has to our knowledge never been described and does not seem ascribable to a systematic bias in the data acquisition. It is illustrated in Figure 4 (ten-fold amplification). The oral tip of the pyramid is very subtly curved towards the left, and the pyramid-epiphysis left junction (landmark set {5, 6, 11}) is higher and narrower than its right counterpart (landmark set {7, 8, 9}).

**Figure 4.**
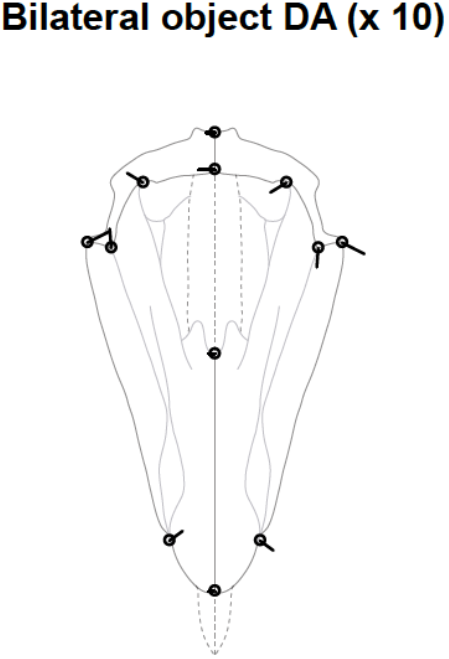
Analysis 2: bilateral object symmetry (C_1*v*_). Bilateral object directional asymmetry (DA) of the pyramids, measured as the difference between the averages of original and reflected configurations (ten-fold amplification).

**Table 2.** Procrustes ANOVA and MANOVA of the pyramid’s bilateral object symmetry only (analysis 2). Abbreviations as in Table 1.

Procrustes ANOVA
| **Effect** | **Conv. df** | **Shape df** | **SS** | **MS** | **F** | **P (param)** | **P(permut)** | **Pillai's trace** | **P(param)** | **P(permut)** |
| --- | --- | --- | --- | --- | --- | --- | --- | --- | --- | --- |
| Pyramid | 4 | 40 | 0.000429 | 0.000011 | 3.312 | <0.0001 | <0.0001 | 1.9941 | <0.0001 | <0.0001 * |
| Reflection | 1 | 10 | 0.010059 | 0.001006 | 365.744 | <0.0001 | <0.0001 | 0.9947 | <0.0001 | <0.0001 |
| Pyramid*Reflection | 4 | 40 | 0.000605 | 0.000015 | 5.497 | <0.0001 | <0.0001 | 1.6941 | <0.0001 | <0.0001 |
| Symmetric error | 45 | 450 | 0.001457 | 0.000003 |  |  |  |  |  |  |
| Asymmetric error | 45 | 450 | 0.001238 | 0.000003 |  |  |  |  |  |  |
| Total error | 45 | 900 | 0.002695 | 0.000003 |  |  |  |  |  |  |

^*^ possible in this design because tested against (symmetric) error (2-way model with only fixed effects)

Among-pyramid variation is mostly marked by the relative narrowing of the foramen magnum, while within-pyramid variation (FA) is spread more evenly across the pyramid and affect the relative length of the hemipyramids (Figure 5).

**Figure 5.**
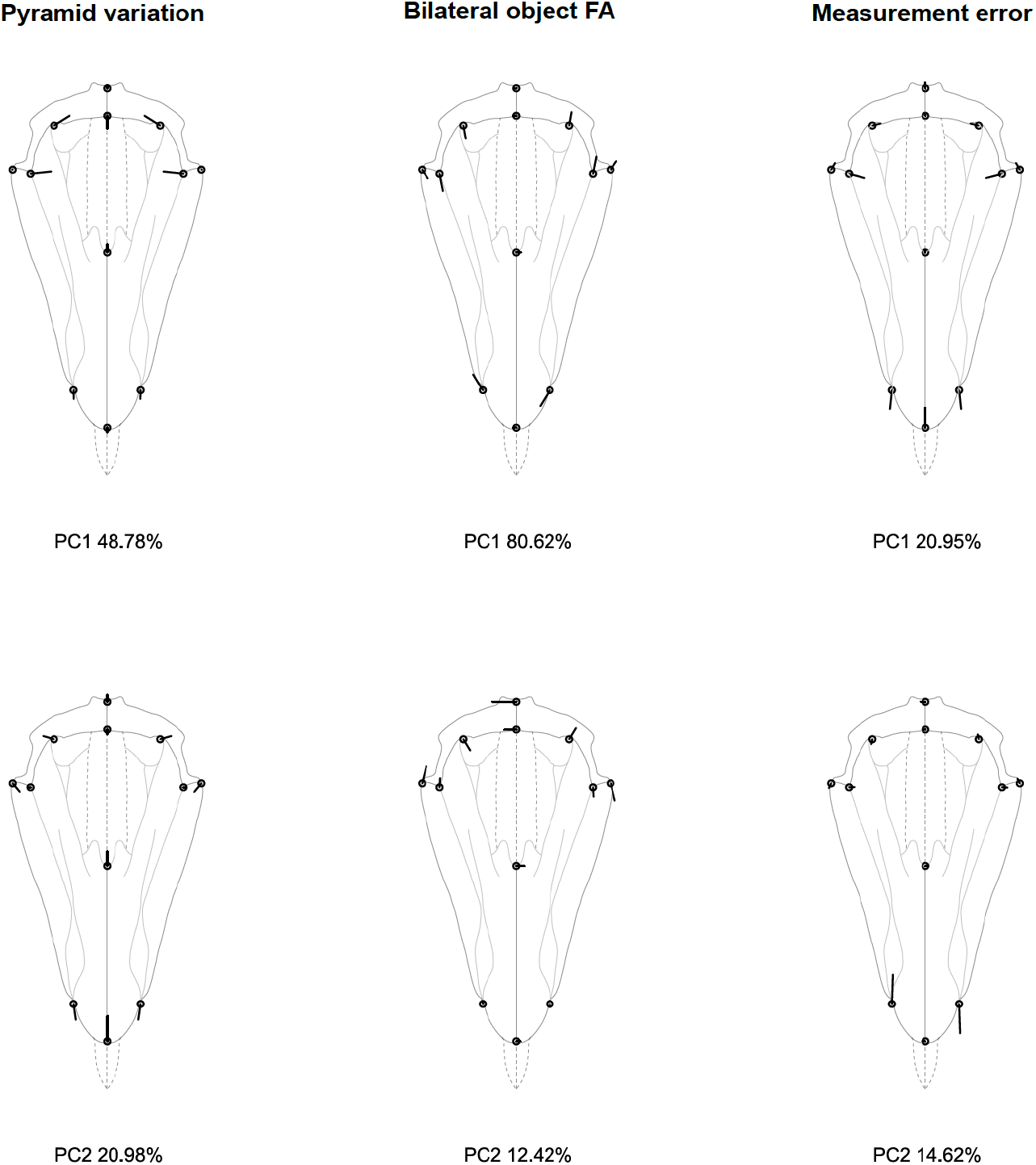
Analysis 2: bilateral object symmetry (C_1*v*_). Principal components describing the patterns of pyramid symmetric variation, bilateral FA, and measurement error. Legend as in Figure 3. Note that for the measurement error panels, the symmetry or asymmetry of the residual vectors with respect to the plane of symmetry depend on the way the symmetric and asymmetric components of the total error variation are distributed along principal components. Here PC1 captures a part of the symmetric variation while PC2 captures an asymmetric part.

### Analysis 3: Bilateral object symmetry nested within rotational matching symmetry of order five (symmetry group C_*5v*_)

The previous analyses were solely concerned with either the bilateral object symmetry of the pyramids or their rotational matching symmetry. Both cases are simplifications of the true complexity of the lantern, since the latter in fact possesses an intertwined combination of the two types of symmetry. We now describe a design that considers the variation in bilateral (left-right) object symmetry at the pyramid level as hierarchically nested within the variation in rotational matching symmetry at the lantern level (symmetry group C_5*v*_, Figure 2c). The design changes into a mixed model ANOVA with nested and crossed factors:

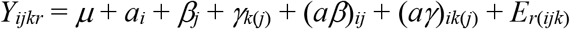

Where *Y_ijkr_* is the measurement for lantern i, pyramid *j*, hemipyramid (or reflection) *k*, replicate *r* (*i* = 1, 2,., 10; *j* = 1, 2,…, 5; *k* =1, 2; *r* =1,2). The parameter *μ* is the grand mean. The main effect *β_j_* is the variation among lanterns ~ *N (0, σ1)*. The main effect *β_j_* is the ‘pyramid’ effect and is the measure of rotational DA. The nested fixed effect *γ_k(j)_* is the ‘hemipyramid’ (or ‘reflection’) effect standing for the nested bilateral DA. The interaction (*αβ*)_*ij*_ is a measure of the lantern rotational FA, ~ *N (0, α;2)*. The interaction *(αγ)_ik(j)_* is a measure of the pyramid bilateral FA, ~ *N (0, σ3)*. *E_r(ijk)_* is measurement error (ME) ~ *N (0, σ4)*.

In this nested design, the among-lantern variation is significant and similar to that of analysis 1, but rotational FA no longer appears significant (Table 3). Bilateral DA is still significant and fairly pronounced within this nested design, with a pattern identical as that shown in Figure 4. Bilateral FA also remains significant but slightly differs from the pattern of shape variation highlighted in analysis 2. Here, the FA variation in the basal part of the pyramid (landmarks {5, 6, 7, 8, 9, 11}) does not appear as strongly correlated with variation at its tip (landmarks {1, 2, 3}) as previously suggested by the integrated pattern of analysis 2, where most of the FA variation was taken up by the first principal component (Figure 6). This suggests some degree of developmental modularity in the architecture of the pyramids.

**Figure 6.**
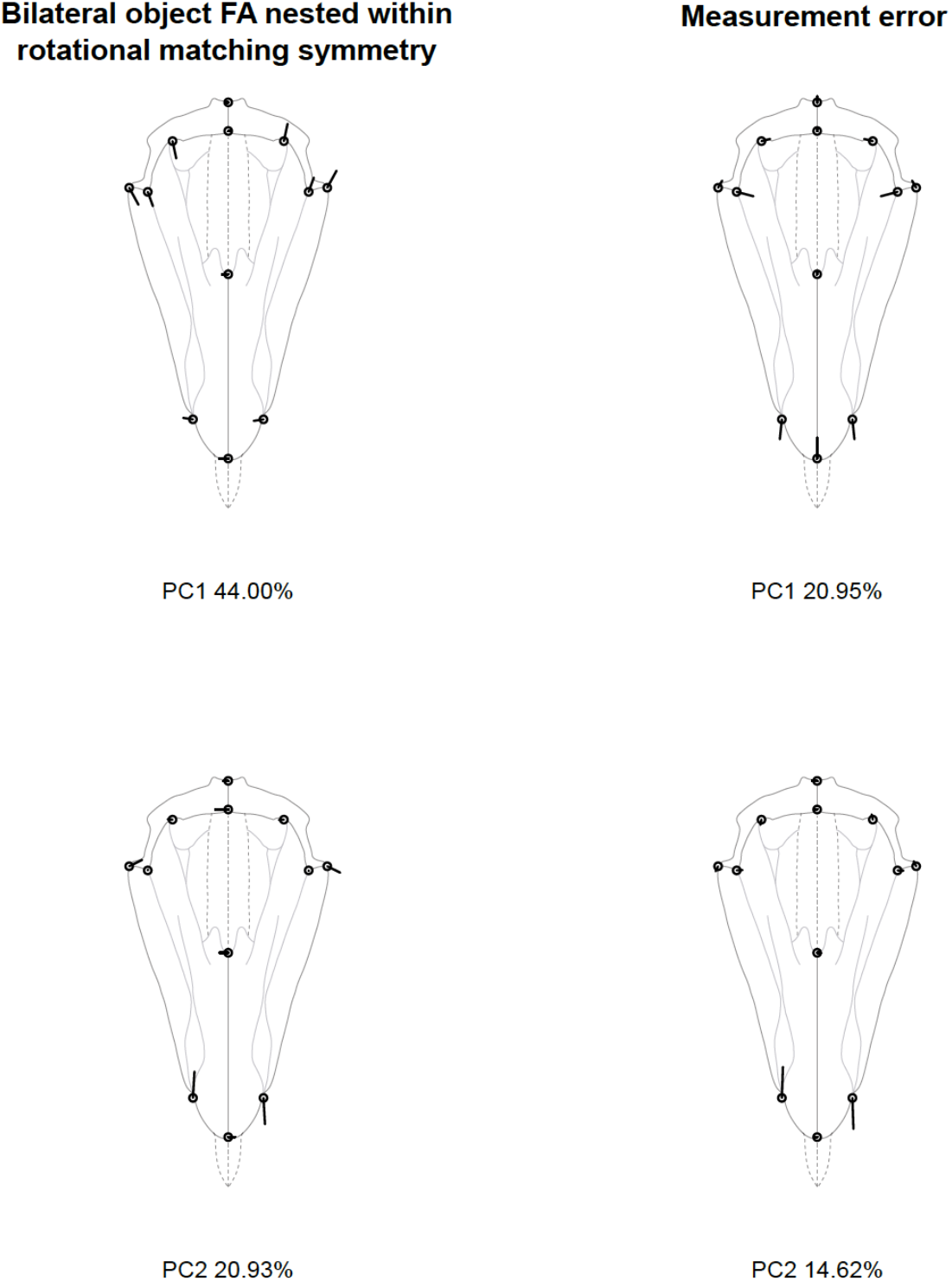
Analysis 3: bilateral object symmetry nested within rotational matching symmetry of order five (C_5*v*_). Principal components describing the patterns of variation for nested bilateral object FA, and measurement error. The patterns of lantern variation and rotational FA are not shown here since they are similar to those shown in Figure 3. Legend as in Figure 3.

**Table 3.** Procrustes ANOVA and MANOVA of the pyramid’s bilateral object symmetry nested within five-fold rotational matching symmetry of the lantern (analysis 3). Abbreviations as in Table 1.

Procrustes ANOVA
| Effect | Conv. df | Shape df | SS | MS | F | P (param) | P (permut) | Pillai's trace | P(param) | P(permut) |
| --- | --- | --- | --- | --- | --- | --- | --- | --- | --- | --- |
| Lantern | 9 | 90 | 0.111274 | 0.001236 | 125.648 | <0.0001 | <0.0001 | 6.8480 | <0.0001 | <0.0001 |
| Pyramid | 4 | 40 | 0.000429 | 0.000011 | 1.090 | 0.33333 | 0.3412 | 1.0869 | 0.3148 | 0.3036 |
| Reflection(Pyramid) | 5 | 50 | 0.010664 | 0.000213 | 12.511 | <0.0001 | <0.0001 | 1.4737 | 0.0071 | 0.0018 |
| Lantern*Pyramid | 36 | 360 | 0.003542 | 0.000010 | 0.577 | >0.9999 | >0.9999 | — | — | — |
| Lantern*Reflection(Pyramid) | 45 | 450 | 0.007671 | 0.000017 | 6.198 | <0.0001 | <0.0001 | 7.6644 | <0.0001 | <0.0001 |
| Symmetric error | 45 | 450 | 0.001457 | 0.000003 |  |  |  |  |  |  |
| Asymmetric error | 45 | 450 | 0.001238 | 0.000003 |  |  |  |  |  |  |
| Total error | 45 | 900 | 0.002695 | 0.000003 |  |  |  |  |  |  |

## Discussion

In this paper, we have discussed the analysis of symmetric and asymmetric shape variation in biological structures exhibiting patterns of hierarchically organised symmetries. We illustrated the problem with the case of the Aristotle’s lantern, a complex organ displaying bilateral symmetry nested within rotational symmetry of order five. Different statistical models could be defined depending on prior hypotheses about the developmental architecture of the lantern and the fundamental unit that was assumed to be repeated to generate the symmetry of the structure. These various designs implied extending to *k* “sides” the standard two-way Procrustes ANOVA for the statistical testing of among- and within-individual variation, dealing with instances of both object symmetry and matching symmetry, and incorporating their nested arrangement for the comprehensive treatment of the lantern’s architecture.

Even if based on a relatively small dataset, the univariate and multivariate approaches and their associated parametric and non-parametric tests all lead to congruent results. Overall, these results outline the importance of considering all levels of symmetric organisation when trying to decipher patterns of individual variation, directional asymmetry and fluctuating asymmetry. While reduced models can detect some effects, those might also be exaggerated or hidden by the treatment of the data implied by a particular model. The full, nested model reveals the absence of rotational DA. This suggests the action of functional constraints ensuring the maintenance of the five-fold rotational symmetry. The feeding efficiency of the lantern is indeed driven by interdependencies among many skeletal pieces and muscles, which might be optimized by the phenotypic similarity of these elements across sides. Likewise, rotational FA is also absent, suggesting high levels of precision in the developmental processes involved in the morphogenesis of pyramids, further supporting the importance of rotational symmetry. Ground-samplers inspired from the Aristotle’s lantern (and its keeled teeth in particular) also base the efficiency of their opening and closing mechanisms on the five-fold repetition of a specific structure akin to a “target phenotype” (Frank et al. 2016). Results differ at the lower level of organisation. Bilateral DA is present and corresponds to a systematic torsion of pyramids, around which can also be detected a significant but subtle degree of bilateral FA. Elucidating the genetic or developmental origin of this DA requires more work, but inadequate analysis of the lantern symmetry would have prevented its detection. Hence, higher order (rotational) symmetry appears to display greater developmental stability than lower order (bilateral) symmetry, but there are no benchmarks in the literature to conclude about the generality of this pattern. One possible cause for these differences could be linked to the spatial pattern of the lantern formation. While the hierarchical structuring might convey the idea of a temporal sequence for the layout of the lower and higher order of symmetry, all the skeletal elements of the lantern form in fact synchronously in the metamorphosing pluteus larva from tri-radiate spicules (Devanesen 1922; Gordon 1926). Nevertheless, the individual pairs of left and right hemipyramids which ultimately fuse to produce the pyramids (bilateral symmetry) are spatially close neighbours during their growth. They are therefore more likely to be prone to feedback interactions generating left-right asymmetry (e.g., competition between growing hemipyramids and interactions with the growing tooth in between them, Klingenberg and Nijhout 1998). The five pyramids themselves are spatially more isolated from each other in the early stage of their formation and might therefore be more immune to such interactions.

We expect that the approach advocated in this paper will be useful in a wide variety of biological contexts and for a broad range of organisms where symmetry is a salient feature of phenotypes. The study of serial homologs in arthropods and vertebrates for instance can be similarly analysed at their different organisational levels as bilateral object symmetry nested within translational matching symmetry (repetition of bilaterally symmetric modules along the body axis). Symmetry in plants is also widely studied (Damerval et al. 2017), and many inflorescences and floral corollae display hierarchical patterns of symmetry (e.g., bilaterally symmetric petals arranged in a rotational fashion). Combined with developmental genetic data, morphometric analysis of patterns of individual variation and asymmetry across levels can refine our understanding of the evolution of symmetry and the identification of its driving factors (e.g., plant-pollinator interactions).

Practically, size and shape data can be obtained from various free software and R packages such as MorphoJ (Klingenberg 2011), tpsRelw (Rohlf 2015), geomorph (Adams and Otárola-Castillo 2013), or shapes (Dryden 2017). The appropriate ANOVA design for complex and/or nested symmetric structures can be implemented with the R function lm() (library {stats}), but the investigator has to carefully adjust the degrees of freedom by considering the shape dimensions of the spaces involved (Goodall 1991; Savriama and Klingenberg 2011). The matrices of sum of squares and cross-products (SSCP) associated with the various factors and interaction terms can be calculated from the effect components retrieved from the lm() outputs using the crossprod() function ({base}). The covariance matrice attached to each effect can then be obtained by subtracting these SSCPs (once divided by their degrees of freedom) following the expected mean squares implied by the ANOVA design (e.g., Klingenberg and McIntyre 1998). These covariance matrices can be decomposed into their eigenvectors and eigenvalues or used as input in MorphoJ to visualize the patterns of individual shape variation, fluctuating asymmetry, and measurement error.

## Conclusion

Geometric morphometric methods are now routinely applied to study bilateral symmetry, but complex types of symmetry also have recently received increasing interest and the morphometric toolkit has expanded accordingly. Here, we have further extended these approaches and suggested a framework for the effective quantification of phenotypic variation and developmental instability in organisms with hierarchically structured symmetries. This allows the partitioning of biological variation into meaningful components of symmetric and asymmetric shape variation at different levels and the analysis of the patterns associated with each of these components. The ability to distinguish components of symmetric variation and asymmetry is of interest in many research contexts including developmental and evolutionary biology, ecology and taxonomic studies. Symmetry on its own has also long been recognized as a fundamental feature of body plan organisation and in the diversification of many major biological groups (e.g., echinoderms, flowering plants, arthropods). The approaches described above offer a unified framework for the characterisation of symmetry and the study of its role in phenotypic organisation, variation, and evolution.

## Acknowledgements

The authors thank the Academy of Finland and the UMR CNRS 6282 “Biogéosciences” for financial support, and Leif Stige, Bruno David, Thérèse Choné, Paul Alibert, and Christophe Lambert for comments and discussions on previous drafts of the manuscript.

## Authors’ contributions

YS and SG conceived the ideas and designed the analyses; YS collected the data; SG wrote the R scripts; YS and SG analysed the data; YS and SG wrote the manuscript. All authors contributed critically to the drafts and gave final approval for publication.

## Data Accessibility

Landmark data will be archived in Dryad [citation to follow when available].

